# Competition and Compromise between Exogenous Probiotics and Native Microbiota

**DOI:** 10.1101/2024.08.15.608074

**Authors:** Zhe Han, Zheng Sun, Qian Zhao, Lingwei Du, Dongyu Zhen, Xinlei Liu, Shuaiming Jiang, Yang-Yu Liu, Jiachao Zhang

## Abstract

Probiotic interventions are effective strategies to modulate the gut microbiome, but how exogenous probiotics compete with native gut microbiota remains elusive. Here, we use a mouse model and a well-documented probiotic *Bifidobacterium animalis* subsp*. lactis* V9 (BV9) to mechanistically investigate its competitive strategies. We perform metagenomic and whole genome sequencing of stool samples and isolated BV9, longitudinally collected from 24 mice orally administered with BV9 and different diets. Results show that a high-fiber diet most effectively supports the colonization of BV9, where BV9 selectively competes with *Parabacteroides distasonis* (*P. distasonis*), rather than extensively with other gut bacteria. By comparing the genomic structures of BV9 and *P. distasonis* isolated during the washout period, we infer their co-evolution mechanisms, highlighting their competition and compromise in utilizing inulin-derived glucose. Finally, our *in vitro* co-culture experiments validate such competitive dynamics. This study fills a critical gap in our understanding of niche competition in colonization.

## INTRODUCTION

Probiotic interventions are effective strategies currently available for modulating the gut microbiome^1,2^. Previous research has extensively evidenced the capacity of probiotics to enhance therapeutic outcomes^3,4^, improve disease states, and regulate physiological processes^5–7^. For example, probiotics have been shown to influence metabolic pathways, immune responses, and even the neuroendocrine system, underscoring their potential as an effective approach to health management^5,8^. The ongoing integration of probiotics into therapeutic regimens emphasizes the need for a deeper understanding of their mechanisms of action^9,10^. This includes not only the identification of beneficial microbial strains but also a comprehension of the interactions between the exogenous probiotic and the native gut microbiota^11^. Even though probiotics with strong tolerances to gastric acids enter the intestinal environment, they must compete for ecological niches within the gut microbiota to colonize effectively^12^. This colonization is not only about survival but involves integrating into the existing microbial community, where they undergo further growth and engage in metabolic activities^13^.

How probiotics compete for ecological niches with the native gut microbiota remains elusive. For instance, depending on the functional overlap between the exogenous probiotics and the native microbiota, probiotics may engage in extensive competition with many native species or relatively selective competition with only a few. Moreover, the competition strength may strongly depend on the availability of common resources, which is still poorly understood. Additionally, the competition may lead to the elimination of one competitor or a compromised balance where all competitors coexist. In the former scenario, the mechanisms of competition between the probiotic and the native gut microbiota are crucial, while in the latter scenario, the nature of their compromises is important. The molecular biological mechanisms driving the competition and subsequent co-evolution are intriguing.

In this work, we utilized a genetically characterized probiotic strain, *Bifidobacterium animalis* subsp*. lactis* V9 (BV9), and a mouse model to address the above questions. Given the significant influence of diet on the structure and function of the gut microbiota^14^, we first established various dietary patterns in mice and discovered that a high-fiber diet best promotes the colonization of BV9. By analyzing the co-occurrence networks of BV9 and native gut microbes under different dietary patterns, we found that BV9 selectively competed with a native species *Parabacteroides distasonis* (*P. distasonis*) for ecological niches. Moreover, by comparing the genome structures of BV9 and *P. distasonis* isolated during the washout period, we elucidated their potential co-evolutionary mechanisms at the genetic level. Finally, subsequent *in vitro* co-culture experiments validated the changes in the competitive relationship between BV9 and *P. distasonis* after co-evolution.

## RESULTS

### Experimental design

The whole experiment lasted seven weeks. The first two weeks served for the establishment of the native gut microbiota associated with a specific dietary pattern. The placebo or probiotic intervention occurred during the third week. The remaining four weeks served as the washout period (**Fig.1a**). We divided 24 C57 mice into four groups, each consisting of six mice. The first group served as the Control group, which was fed with Normal Diet throughout the experiment and received daily oral administration of placebo (300ul skim milk) during the third week. The other three groups were fed with three different diets: Normal Diet (ND), High-Fat Diet (HFAD), and High-Fiber Diet (HFID), respectively, throughout the experiment, and received daily oral administration of 2×10^9^ cfu BV9 along with 300ul of skim milk during the third week. We designated the last day of the placebo or BV9 gavage as Day 0, and collected fecal samples on days 1, 7, 14, 21, and 28 during the washout period. In total, we collected 120 fecal samples (5 time points * 4 groups * 6 mice), which were then subjected to metagenomic sequencing to investigate the response of the gut microbiome shaped by different dietary patterns to probiotic administration.

To assess the colonization outcome of BV9 and explore the genetic alterations of BV9 post-colonization, we isolated and cultured BV9 strains daily from fecal samples during the washout period. The strains identified as BV9 via 16S rRNA gene sequencing were further examined using Whole-Genome Sequencing (WGS) to investigate their genetic variations (**Fig. 1a**). We used isolated strains and the Whole-Metagenome Sequencing (WMS) data to ascertain the colonization dynamics of BV9 and the dynamics of the gut microbiome during the washout period. Employing the co-occurrence network analysis, we identified a potential selective competition between BV9 and *P. distasonis* (**Fig. 1b**). Subsequently, with a hypothesis that inheritable mutations (e.g., SNVs) significantly contribute to such adaptive evolution, we explored the co-evolutionary mechanisms between BV9 and *P. distasonis* at the genetic level using the WGS data and validated our findings through *in vitro* co-culture experiments (**Fig. 1c**).

**Figure 1.**
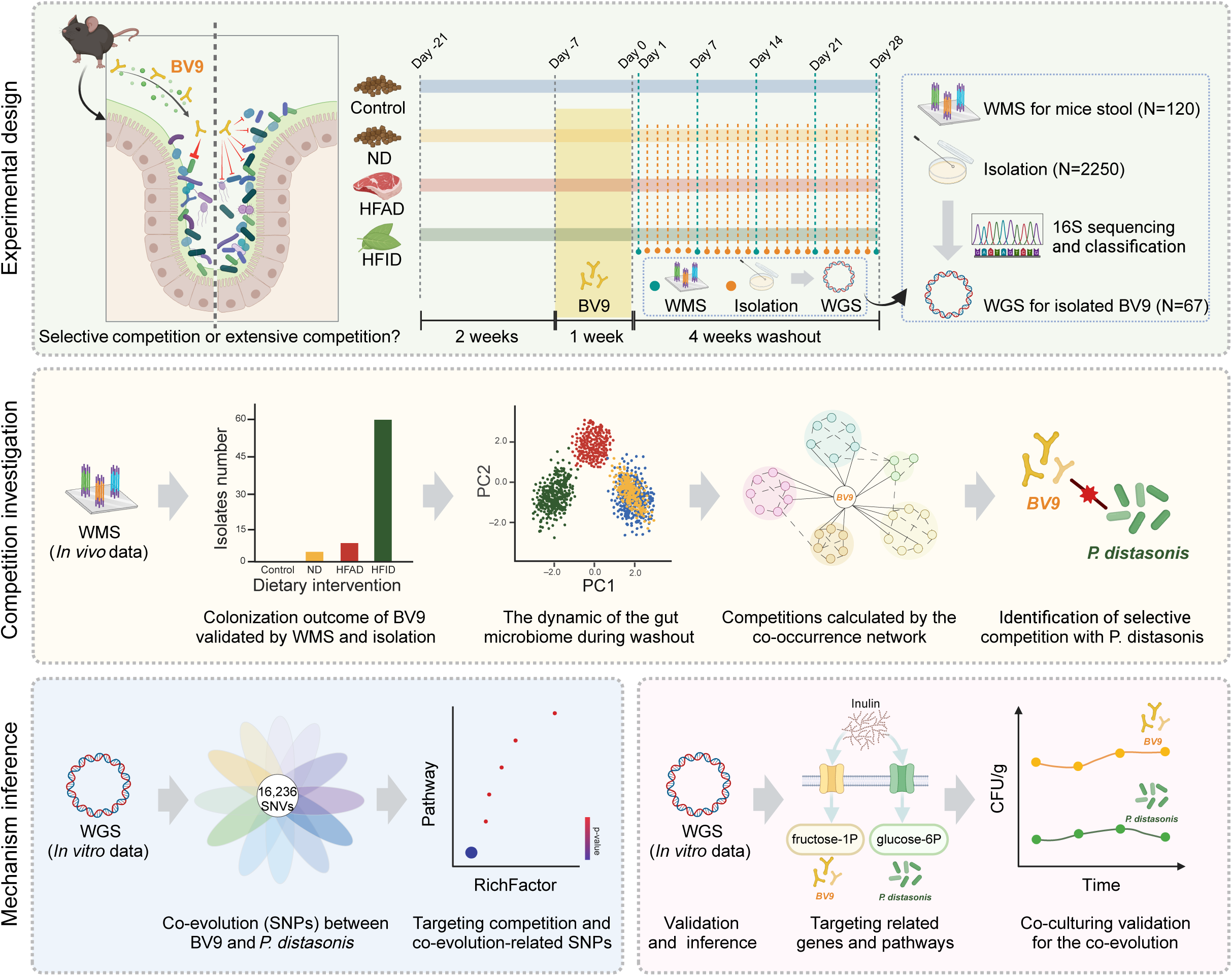
Conceptual framework of the study. (**a**) To investigate the impact of probiotics on the native gut microbiota and their competitive interactions, we selected BV9 as the study subjects and used C57 mice as the experimental model (**Methods**). Initially, the mice underwent a two-week dietary intervention with a normal diet (ND), high-fiber diet (HFID), and high-fat diet (HFAD). Subsequently, all groups except the control were gavaged with BV9 for one week. Stool samples were collected on days 1, 7, 14, 21, and 28 after cessation of probiotic gavage, resulting in a total of 120 fecal samples for metagenomic sequencing to study the response of gut microbiota to probiotic intervention under different dietary conditions. Additionally, BV9 strains were also isolated and cultured daily from the fecal samples. Preliminary screening (by using PCR) identified 2250 suspected BV9 isolates and 16S rRNA sequencing confirmed 67 of these as BV9. Then, Whole-genome re-sequencing (WGS) was then performed to further analyze the genetic variations of these strains. (**b**) First, we evaluated the colonization efficiency of BV9 under different dietary patterns by combining WMS data and BV9 isolation verification results. Next, we analyzed the dynamic changes in gut microbiota during the washout period using WMS data. Subsequently, we constructed co-occurrence networks for each dietary pattern, identifying *P.distasonis* as a species competing with BV9 within the native gut microbiota. (**c**) Based on WGS data from isolated strains, we identified SNVs closely associated with colonization in both BV9 and *P.distasonis*. We then performed pathway differential analysis on the genes and pathways containing these SNVs to pinpoint genes and pathways related to competition, uncovering the molecular biological mechanisms of co-evolution. Finally, we co-cultured these two strains to validate the co-evolutionary changes.

### Impact of Dietary Patterns on BV9 Colonization

To assess the colonization outcome of BV9, we isolated a total of 2,250 bacterial isolations from all mice during the four-week washout period. Following verification with 16S rRNA gene sequencing and confirmation via WGS (**Methods**), we identified in total 67 isolations as BV9. Upon comparing the number of BV9 isolations across the three groups, we consistently identified more BV9 isolations from the HFID group throughout the washout period, totaling 54 isolations. In contrast, only 13 BV9 isolations were obtained from the HFAD and ND groups combined during the washout period (**Fig. 2a**).

**Figure 2.**
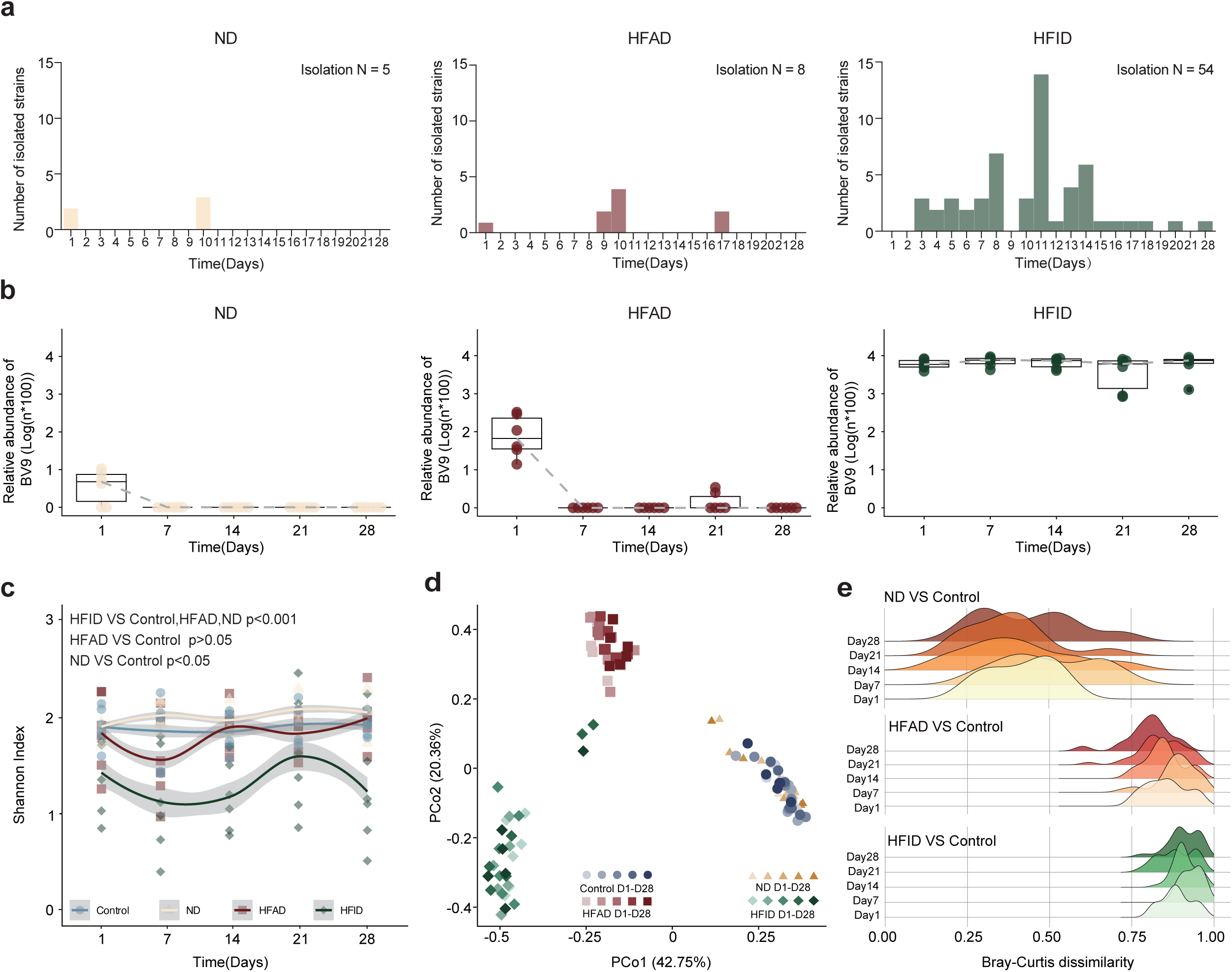
Colonization of BV9 and the dynamic of the mice gut microbiota under different dietary conditions. (**a**) The number of BV9 isolates obtained from fecal samples during the washout period under different dietary conditions. (**b**) Comparison of the relative abundance of BV9 during the washout period under different dietary conditions. The gray dashed lines in the box plots connect the medians at each time point. (**c**) The Shannon index during the washout period in different dietary groups. (**d**) Principal Coordinate Analysis (PCoA) based on Bray-Curtis dissimilarity. It illustrates the dynamic of the gut microbiota under different dietary conditions during the washout period. Each point represents a sample, color-coded by diet type and labeled by time point (days 1, 7, 14, 21, 28). (**e**) Ridge plots showing the distribution of Bray-Curtis dissimilarity between the mice in the control group and the mice in other dietary groups.

To further validate the colonization efficiency of BV9, we profiled the WMS data and compared the relative abundance of *Bifidobacterium animalis* among the groups. This comparison was conducted at the species level because BV9 was the only strain of *B. animalis* observed in all mice. The results from the WMS data aligned with those obtained from the isolations, demonstrating a consistently higher abundance of BV9 in the HFID group with respect to the other groups (**Fig. 2b**). Thus, we conclude that a high-fiber diet facilitates the colonization of BV9, whereas a high-fat diet or normal diet does not. This agrees well with a previous study that high-fiber diet can benefit the colonization of *Bifidobacterium*^15^.

### Impact of Probiotic Intervention on the Gut Microbiota

To investigate the transient and long-term impact of probiotic intervention, we compared the dynamics of the gut microbiome in mice under different dietary patterns during the washout period (**Methods**). We first performed the alpha- and beta-diversity analyses. For the alpha diversity, although a decrease followed by an increase in the Shannon index was observed in the HFAD group, overall, we found no significant change of the Shannon index for the HFAD group, with respect to the Control group. This is consistent with the observation that BV9 was constrained in colonization in the HFAD groups, and may only have a transient impact on the native microbiota. In contrast, the HFID group displayed a significantly lower Shannon index with respect to other groups (**Fig. 2c**). This is because BV9 colonization was facilitated in the HFID group, where BV9 eventually predominated the mice gut microbiota with an average relative abundance 0.62.

Regarding beta diversity, we performed the Principal Coordinates Analysis (PCoA) using the species-level Bray-Curtis dissimilarity measure. We found that the gut microbiomes of the HFAD and HFID groups differ significantly from the ND and Control groups, throughout the washout period (**Fig. 2d, Methods**). This finding implies that diet remains a primary factor shaping the structure of the gut microbiome, which is consistent with previous studies^16–18^. Notably, the HFID group showed the most substantial change in gut microbiome structure after the probiotic intervention compared to the Control group (**Fig. 2e**). Further, we examined changes in the gut microbiomes along the time within each group following probiotic intervention, and we found that while the changes in the HFAD group were most convergent (e.g., at each time point, the Bray-Curtis dissimilarities between different individuals in the same group are greater in HFAD than other groups.), HFID were most divergent (Fig. S1).

We next performed differential abundance analysis at both species and functional pathway levels between each of the three diet groups and the Control group across all the five time points during the washout period (Fig. S2, **Methods**). We found that the gut microbiome of the HFID group with long-term BV9 colonization displayed the most significant differences at both species and pathway levels after the probiotic intervention (36 differentially abundant species compared to 33 for the HFAD group or 14 for the ND group, and 50 differentially abundant pathways compared to 36 for the HFAD group or 10 for the ND group; Fig. S2a, S2b).

At the species level, we observed significant differences between each of the three diet groups and the Control group. For the ND group, *Lactobacillus reuteri* (Day 21,28; p-value = 0.0223, 0.0029; average log2 ratio = 3.2439), *Lactobacillus johnsonii* (Day 14; p-value = 0.0068; log2 ratio = 1.9832), and *Akkermansia muciniphila* (Day1,7,14; p-value = 0.0053, 0.0237, 0.0291; log2 ratio = 2.0609) are representative differentially abundant species among the 14 differentially abundant species. For the HFAD group, *Bacteroides vulgatus* (p-value < 0.05 for all time points, log2 ratio = 11.2439), *Lachnospiraceae bacterium 3 2* (Day1, 7, 28; p-value = 0.003, 0.04, 0.02; log2 ratio = 4.8665), and *Bacteroides faecichinchillae* (log2 ratio = −6.8220) are representative differentially abundant species among the 33 differentially abundant species. For the HFID group, the greatest differences were observed for *Muribaculaceae bacterium DSM 103720* (log2 ratio = −7.0383), *Parasutterella excrementihominis* (log2 ratio = 5.2224), and *Alistipes shahii* (log2 ratio = −3.0606) among the 36 differentially abundant species.

At the pathway level (**Methods**), the ND group showed the smallest changes, with only minor differences in 10 pathways, such as L-lysine biosynthesis II (Day 7,28; p-value = 0.001, 0.0001, log2 ratio =3.5774) and super pathway of pyrimidine ribonucleotides de novo biosynthesis (Day1,7; p-value = 0.0002, 0.002, log2 ratio = 2.0898). The HFAD group displayed a variation in 36 differential pathways, such as phospholipases (log2 ratio = 6.8051) and stearate biosynthesis II (log2 ratio = 6.0761). The HFID group exhibited the most differences post-probiotic intervention, including significant changes in pathways involved in palmitate biosynthesis (p-value < 0.001 for all time points, average log2 ratio = 11.4160) and petroselinate biosynthesis (log2 ratio = 10.8676) among 50 differentially abundant pathways.

### Selective competition between BV9 and native gut microbiota

To understand the colonization process of BV9, such as its competition for ecological niches with the native gut microbiota, we further investigated the dynamics between various species within the gut microbiota during the washout period under different dietary conditions, aiming to explore the potential interactions between the exogenous probiotic and the native gut microbiota. Firstly, using Spearman correlation coefficients as edges and taxa at the species level as nodes, we constructed co-occurrence networks for each diet group based on microbiome data during the washout period (only nodes with moderate correlation, i.e., R > 0.4, Spearman correlation test p < 0.05 are shown in **Fig. 3a**, **Methods**). We observed that in the HFAD and ND groups, the co-occurrence networks exhibited relatively high density (0.2 and 0.238, respectively), but the number of species significantly associated with BV9 was limited to 2 and 4, respectively. However, in the HFID group, although a lower overall network density (0.096) is observed, the number of species significantly associated with BV9 is the highest, including *P. distasonis*, *Parabacteroides goldsteinii*, *Bacteroides thetaiotaomicron*, and eight others.

**Figure 3.**
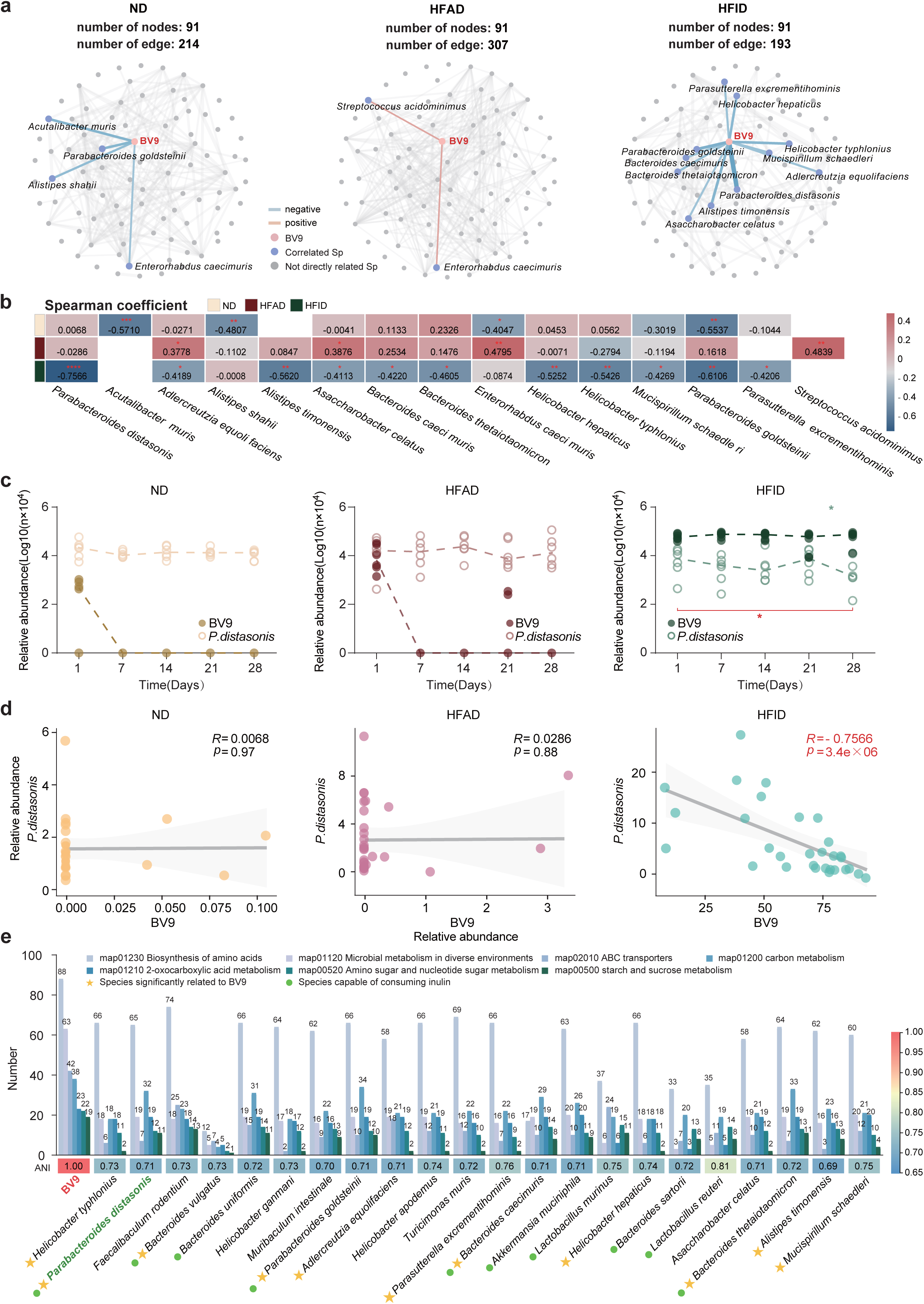
Potential interactions between BV9 and native gut microbiota under different dietary conditions. (**a**) Co-occurrence networks of the gut microbiota were generated by using the species-level taxonomic profiles in ND, HFID, and HFAD groups separately. Nodes represent bacterial species, and the width of the edges indicates the strength of the correlation (Spearman correlation coefficient). Blue edges represent negative correlations, while red edges represent positive correlations. (**b**) The heatmap shows the Spearman correlation coefficients and p-values between BV9 and associated bacterial species under different dietary conditions (*: p < 0.05, **: p < 0.01, ***: p < 0.001). (**c**) Line plots illustrate the change in the relative abundance of BV9 and *P.distasonis* during the washout period in ND, HFID, and HFAD groups. (**d**) The correlation between the relative abundances of BV9 and *P.distasonis* in the ND, HFID, and HFAD groups. R refers to the Spearman correlation coefficient. **(e)** Bar chart illustrates the number of functionally annotated genes within inulin-related carbon metabolism pathways for BV9, its significantly associated strains, and the top 20 species by relative abundance. Heatmap below shows the average nucleotide identity among different bacterial species. Asterisk: Denotes strains significantly associated with BV9. Green dot: Indicates strains capable of consuming inulin.

To further clarify whether BV9’s colonization process involves selective or extensive competition, we compared the Spearman correlation coefficients of species that were significantly associated with BV9 across three groups (highlighted in **Fig. 3a**). We found that *P. distasonis* and BV9 demonstrated the strongest correlation, which only existed in the HFID group where BV9 successfully colonized (R = −0.7566, p = 1.31×10^6^, compared to HFAD R = −0.0286, p > 0.05, and ND R = −0.0068, p > 0.05). Moreover, the correlation between BV9 and *P. distasonis* is much higher than that between BV9 and other bacteria (**Fig. 3b**). Additionally, by comparing the relative abundance of all gut bacteria capable of metabolizing inulin, *P. distasonis* exhibits the highest abundance (Fig. S4). All the above results suggest that BV9 intake may specifically threaten the ecological niche of *P. distasonis*.

To validate this hypothesis, we further compared the change in the relative abundance of all bacteria that are significantly associated with BV9 in the HFID group over time (Day 1 to Day 28) (Fig. S3, **Methods**). We found that only *P. distasonis* showed significant fluctuations in abundance in the HFID group (In the HFID group, D1 vs D28, t-test p = 0.026, **Fig. 3c, 3d**), while BV9’s limited colonization in the HFAD and ND groups did not significantly impact *P. distasonis*’s relative abundance. All these results indicate that BV9 is selectively competing with *P. distasonis* during colonization.

To understand the selection of competitors by BV9, we compared the genome structure and functional capacity between BV9 and all its correlated species, as well as the top 20 species in relative abundance. We hypothesized that BV9 might select its competitors based on more similar genome structure or functional capacity (**Fig. 3e**, **Methods**). By comparing the number of functional genes annotated in inulin-related carbon metabolism pathways, we found that the count between BV9 and *P. distasonis* is not the highest, nor is their genome similarity. This suggests that the selective competition between BV9 and *P. distasonis* might be simply related to its highest relative abundance among all inulin-consuming species^19–23^.

### Exploring the mechanism of the co-evolution between BV9 and P. distasonis

The competition between BV9 and *P. distasonis* did not result in the elimination of *P. distasonis* but a new state where both BV9 and *P. distasonis* stably coexist. We hypothesize that the coexistence of BV9 and *P. distasonis* is associated with heritable genetic alterations. To test this hypothesis, we first investigated single-nucleotide variants (SNVs) in BV9 that are associated with its colonization. We then categorized these SNVs into coding and non-coding variants to explore their functions. For SNVs located in coding regions, we examined whether they are nonsynonymous mutations and if they are under positive selection. For those in non-coding regions, we investigated their potential impact on the expression of downstream genes. We conducted the same analysis for *P. distasonis*. Finally, by comparing the enrichment of downstream genes and their pathways, we aimed to elucidate the genetic level mechanisms of such co-evolution (**Fig. 4a, Methods**).

**Figure 4.**
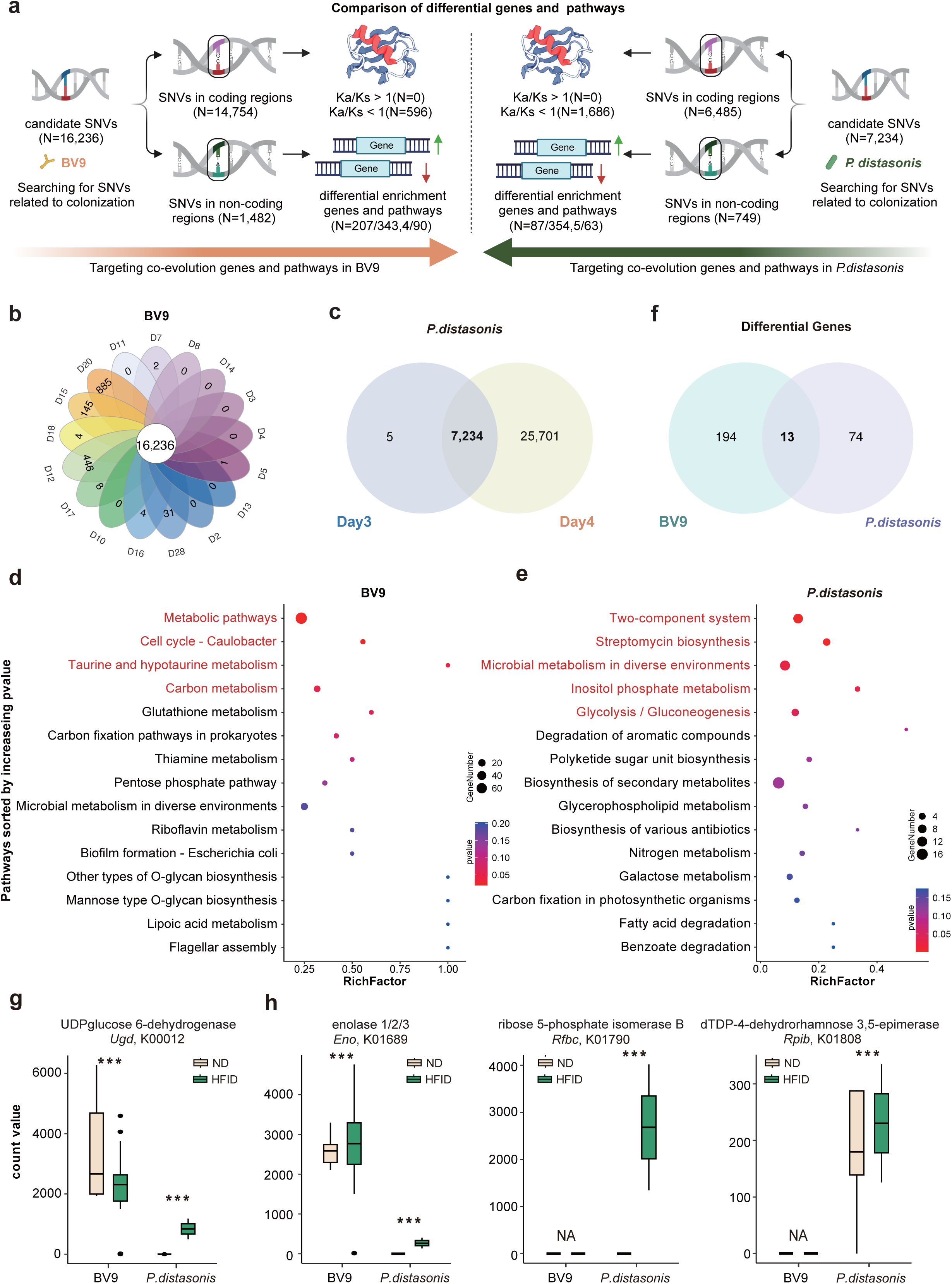
Potential co-evolution mechanism between BV9 and *P. distasonis*. (**a**) Conceptual diagram illustrating our workflow in exploring their co-evolution mechanisms. Candidate SNVs associated with colonization are analyzed differently by their original regions. For SNVs in coding regions, we assess whether they are non-synonymous mutations and if the genes are under positive selection. For SNVs in non-coding regions, we examine their potential impact on downstream gene expression and perform differential enrichment analysis on these genes and pathways to identify the possible gene expression regulation. (**b-c**) Bubble plots shows the results of pathway enrichment analysis for non-coding SNVs in BV9 and *P. distasonis*, with pathways ranked by increasing p-values. In BV9, three pathways are significantly enriched, while in *P. distasonis*, four pathways are significantly enriched (p < 0.05). (**d-e**) Selection of candidate SNVs using the Venn diagram. The intersection is considered to be associated with the colonization of BV9 and *P. distasonis*. (**f**) Venn diagram shows the same differentially expressed genes between BV9 and *P. distasonis*, a total of 13. (**g-h**) Expression levels of differentially expressed genes related to inulin metabolism in both BV9 and *P.distasonis* (*: p < 0.05, **: p < 0.01, ***: p < 0.001).

We first identified SNVs in BV9 associated with its colonization, positing that those SNVs facilitating adaptive adjustments in BV9 should not be exclusive to a single time point but rather persist throughout the entire washout period with high-fiber diet. Given that the presence of other gut bacteria in the WMS data could affect SNV calling, we performed whole-genome sequencing (WGS) on BV9 isolates (n=54) collected at various time points to identify colonization-related SNVs (**Methods**). Based on this premise, we utilized the WGS data to target the intersection of SNVs present at all the BV9-isolation time points (Day 3, 4, 5, 6, 7, 8, 10, 11, 12, 13, 14, 15, 16, 17, 18, 20, 28), uncovering 16,236 shared SNVs across all time points (**Fig. 4b**). Moreover, we conducted identical analyses for *P. distasonis* and identified 7,234 SNVs (Day 3, Day 4; **Fig. 4c**) in total.

For SNVs located in coding regions, we assessed whether selective pressure had acted on the coding genes and led to nonsynonymous mutations. If these genes were not under natural selection, according to the neutral theory of molecular evolution, the rate of nonsynonymous to synonymous substitutions would be equal, i.e., K_a_/K_s_ = 1 (indicating neutral evolution). A ratio of K_a_/K_s_ > 1 would suggest that the gene is under positive selection, while K_a_/K_s_ < 1 indicates purifying selection^24^. We found that among the 14,754 SNVs located in the coding regions of BV9, none exhibited K_a_/K_s_ > 1. Similarly, in *P. distasonis*, out of 6,485 SNVs located in coding regions, none showed K_a_/K_s_ > 1 either (**Fig. 4a**).

For SNVs located in non-coding regions, which typically influence the expression of downstream genes ^25^, we first conducted a differential abundance analysis of these genes, followed by a differential enrichment analysis of the pathways in which they participate (**Methods**). For BV9, 1,482 SNVs were identified in non-coding regions, potentially impacting the regulatory expression of 343 genes, and 207 out of 343 genes showed significant differences among different dietary groups with clear annotation. Further pathway enrichment analysis revealed significant effects on four pathways: Metabolic pathways (p = 0.02), Cell cycle - Caulobacter (p = 0.02), Taurine and hypotaurine metabolism (p = 0.04) and Carbon metabolism (p = 0.05) (**Fig. 4d**). For *P. distasonis*, we identified 749 SNVs located in non-coding regions, potentially affecting the regulatory expression of 354 genes, with 87 genes (without unannotated hypothetical proteins) showing significant differences across different dietary groups. Subsequent pathway enrichment analysis indicated significant impacts on five pathways: Two-component system (p = 0.001), Streptomycin biosynthesis (p = 0.003), Microbial metabolism in diverse environments (p = 0.03), Inositol phosphate metabolism (p = 0.03), and Glycolysis / Gluconeogenesis (p = 0.04) (**Fig. 4e**).

Given that there is no overlap in the pathway level, we further compared the differentially duplicated genes between the two bacteria to further understand the molecular biological mechanisms of the co-evolution between BV9 and *P. distasonis* (**Methods**). Our analysis revealed 13 differential genes shared with BV9 and *P. distasonis* (**Fig. 4f**), and interestingly, we found that only UDP-glucose-6-dehydrogenase (*Ugd*, K00012, BV9: log2fc = −0.3916, *P. distasonis*: log2fc = 12.7571) out of the 13 genes showed an opposite changing trend in BV9 and *P. distasonis*, which is closely related to Glucose 6-phosphate (Glucose-6P) metabolism and carbohydrate metabolism (**Fig. 4g**).

### Co-evolutionary dynamics in inulin metabolism

Inulin is metabolized by bacteria primarily via two pathways^26,27^: (1) The first involves the breakdown of inulin into fructose units, which are converted into D-fructose-1-phosphate. This is transported into bacterial cells and enters the glycolysis pathway as D-fructose 1,6-bisphosphate, facilitating energy production through carbohydrate metabolism. (2) The second pathway begins with the transformation of inulin-derived glucose into glucose-6-phosphate. This is then converted into D-Glucose-1-Phosphate and further metabolized into vital compounds such as UDP-glucose, Phosphoenolpyruvate and dTDP-4-keto-6-deoxy-D-glucose, playing crucial roles in biosynthetic and metabolic processes within the bacterial cells and is an important aspect of carbohydrate metabolism^27^.

We next reviewed the differential genes (using a log2fc threshold of 0.4) associated with inulin metabolism and compared their changing mode in both BV9 and *P. distasonis* (**Fig. 5**). For example, in *P. distasonis*, we identified three additional genes associated with inulin metabolism besides *Ugd*, they are enolase 1/2/3 (*Eno*, K01689, log2fc = 18.5834), ribose 5-phosphate isomerase B (*Rpib*, K01808, log2fc = 0.6284), and dTDP-4-dehydrorhamnose 3,5-epimerase (*Rfbc*, K01790, log2fc = 16.8898) (**Fig. 4h)**. However, in BV9, only a significant decrease in *Ugd* (log2fc = −0.3916) was noted. This suggests that *P. distasonis* may have enhanced its use of the second pathway, whereas BV9 appears to have reduced its reliance on this pathway. Therefore, we infer that during colonization, BV9 predominantly utilizes the first pathway (fructose metabolism), which is more efficient in utilizing inulin^28^, while *P. distasonis* has adapted to primarily exploit the second pathway (glucose metabolism). These adaptations indicate a co-evolutionary compromise between the two species in the utilization of inulin.

**Figure 5.**
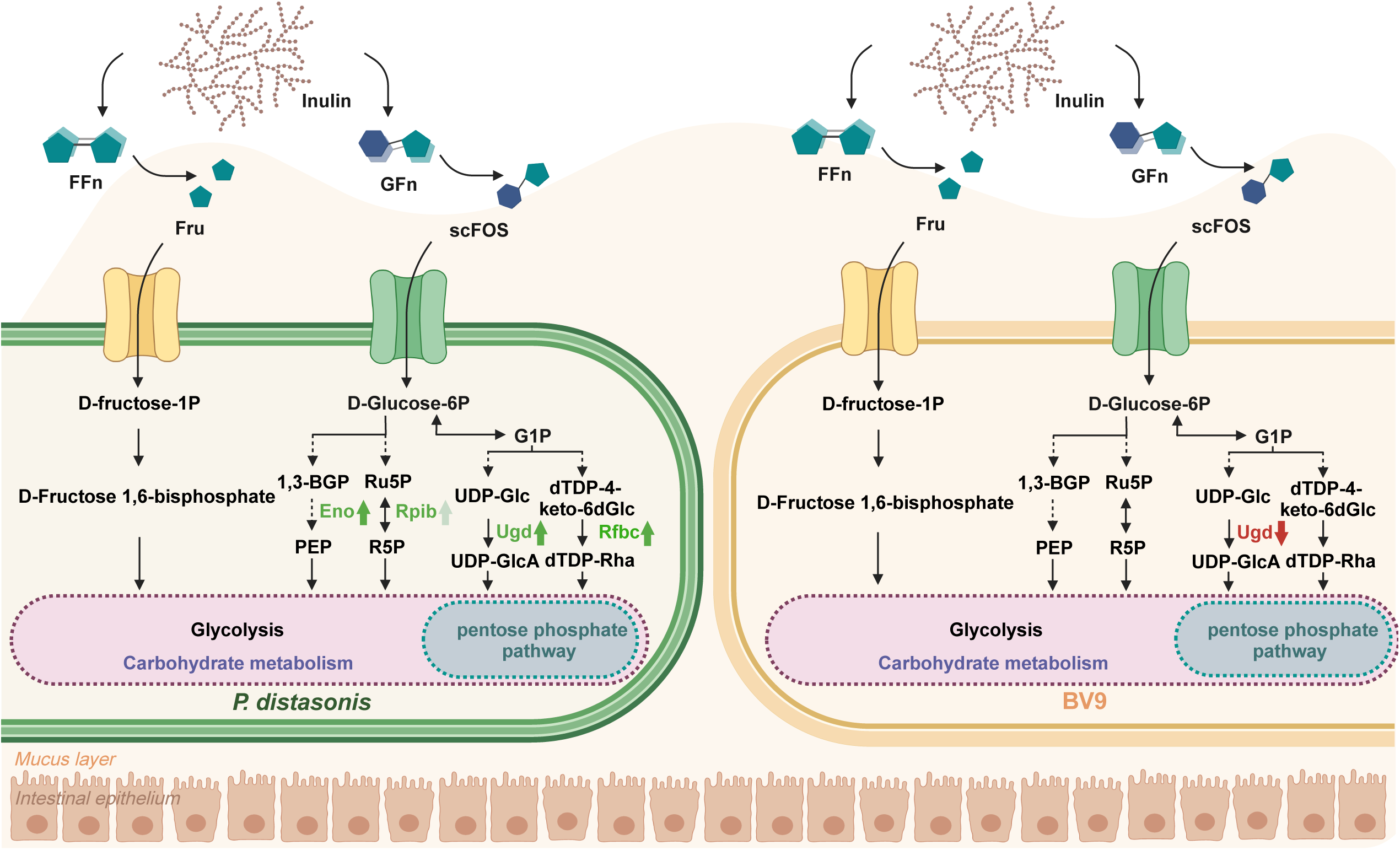
Co-evolutionary mechanisms between BV9 and *P. distasonis* under a high-fiber Diet. Upon entering the gut, inulin is metabolized via two primary pathways. The first pathway degrades inulin into fructose units, generating energy through glycolysis, while the second converts glucose from inulin into essential cellular metabolites like UDP-galactose. Genomic analysis reveals that both BV9 and *P. distasonis* possess genes for fructose metabolism, with no gene duplication observed post-colonization. Notably, *P. distasonis* shows significant enrichment of genes associated with glucose metabolism (*<u>Rpib</u>, Eno, Ugd*, *Rfbc*), indicating enhanced pathway utilization. In contrast, BV9 exhibits a reduction in the *Ugd* gene, suggesting decreased dependency on this pathway. These adaptations may represent a co-evolution strategy where BV9 primarily utilizes the fructose pathway to efficiently metabolize inulin, while *P. distasonis* adapts to predominantly use the glucose pathway. These adaptations suggest a co-evolutionary compromise in inulin utilization between the two bacteria. Dark green arrows represent log2fc>1, and light green arrows represent log2fc<1.

### Changes in competitive modes in co-culture experiment

To further validate the co-evolution between BV9 and *P. distasonis*, we conducted *in vitro* co-cultivation experiments with two sets of BV9 and *P. distasonis* strains: (1) Pre-coevolution: the original strains of BV9 and *P. distasonis*, which had not undergone BV9 colonization in the mouse gut, and (2) Post-coevolution: strains of BV9 and *P. distasonis* isolated on the Day 3 of the washout period after successful colonization of BV9 in the HFID group. Each set was co-cultured under different simulated dietary conditions, including a standard fluid trypticase-phytone-yeast extract (TPY) medium for a normal diet, an inulin-substituted TPY medium replacing glucose with inulin to simulate a high-fiber diet, and a low-carbohydrate medium to simulate a high-fat diet. Over a 48-hour cultivation period, qPCR was performed every two hours and repeated three times (**Methods**).

We observed that in co-cultures of the original BV9 and *P. distasonis* strains (Pre-coevolution), there was always a negative correlation between the growth of BV9 and *P. distasonis* in either standard TPY, low-carbohydrate TPY media, or the inulin-substituted TPY medium (**Fig. 6a-c**), suggesting a significant competitive interaction between BV9 and *P. distasonis* (R = −0.69, p < 0.001, R = −0.81, p < 0.001, R = −0.52, p < 0.01 separately). For the strains isolated after *in vivo* BV9 colonization (Post-coevolution), the competition remains in standard TPY medium or low-carbohydrate media (R = −0.81, p < 0.001, R = −0.75, p < 0.001; **Fig. 6d, 6e**). However, competitive interaction disappeared in the inulin-substituted TPY medium (R = −0.26, p = 0.13; **Fig. 6f**). Collectively, these results indicate that (1) the original BV9 and *P. distasonis* exhibit *in vitro* competitive interactions with any carbon source in media; (2) After adaptation through the *in vivo* BV9 colonization process, the symbiotic relationship between BV9 and *P. distasonis* alters, permitting conditional *in vitro* coexistence only when inulin is the carbon source; (3) The heritability of co-evolutionary changes.

**Figure 6.**
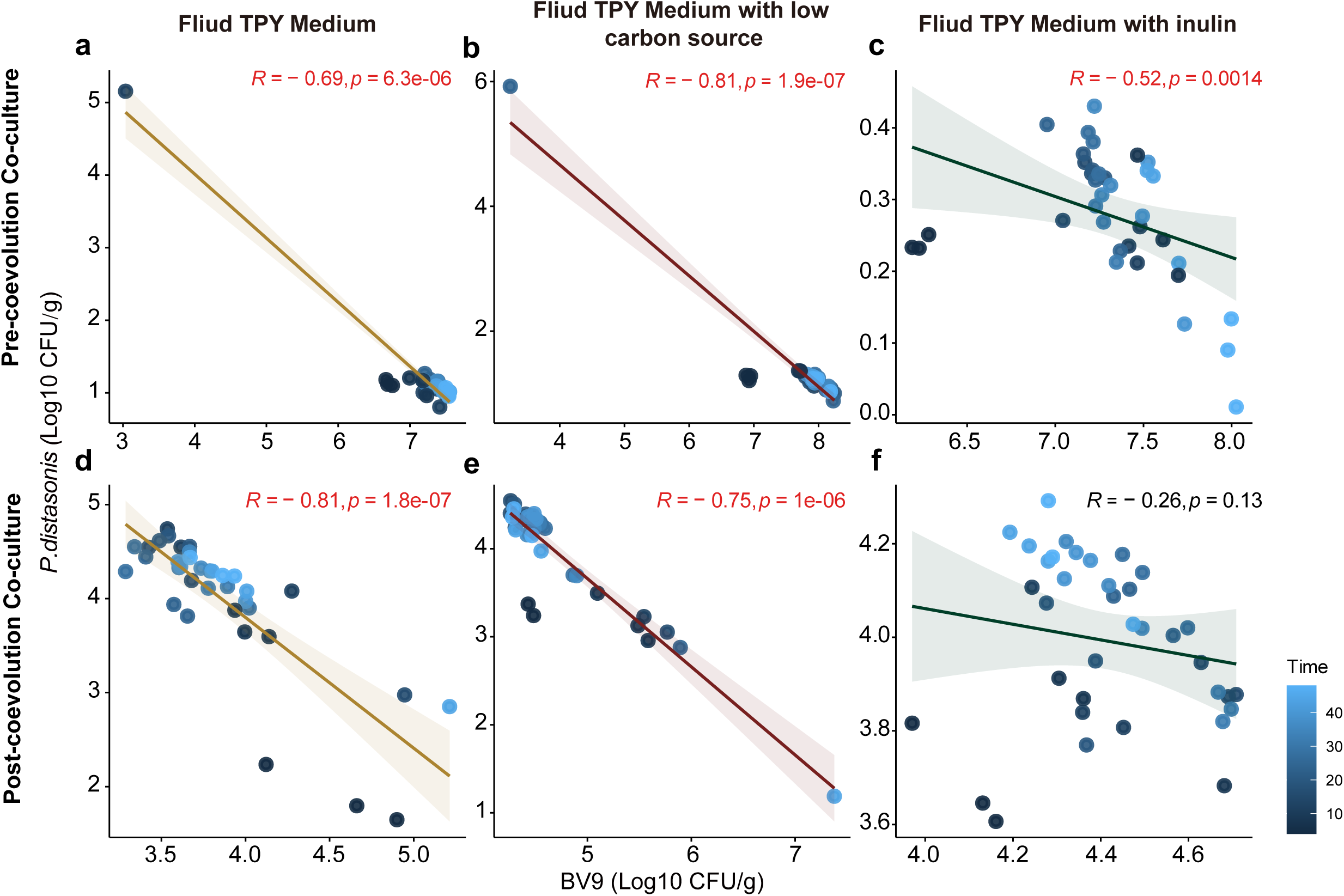
Correlation between the relative abundance of BV9 and *P. distasonis* under different fluid TPY mediums before and after co-evolution. Correlations between the growth of the original strains of BV9 and *P. distasonis* (pre-coevolution) in (**a**) standard fluid TPY medium, (**b**) fluid TPY medium with inulin (to simulate HFID), and (**c**) fluid TPY medium with a low carbon source (to simulate HFAD). (**d-f**) Relationship between isolated strains of BV9 and *P. distasonis* (post-coevolution) under different fluid TPY medium conditions.

## DISCUSSION

Although previous studies have discussed various aspects of how probiotic bacteria interact and adapt within ecological niches ^29–31^, detailed discussions on the selectivity of niche competition during colonization are still lacking. In this paper, we systematically explore the mechanisms of niche competition during the colonization process of the probiotic BV9 in a mouse model, demonstrating its nutrient-dependent and single-species targeting competition strategy. Based on the notably strong negative correlation between their relative abundances, we propose that BV9 undergoes selective competition with *P. distasonis* with inulin available as nutrition. This is based on observations during the washout period rather than the gavage period, as continuous supplementation of BV9 during the gavage could bias the self-driven dynamics of the gut microbiota, thus precluding an objective identification of competitors through co-occurrence networks. Interestingly, although *P. distasonis* doesn’t display the highest genomic similarity and functional overlap among the native species, it exhibits the highest relative abundance among all inulin-consuming species. This explains why BV9 selectively competes with *P. distasonis*. Additionally, we examined genomic changes (in terms of SNVs) in all species significantly associated with BV9 using the WMS data. By analyzing mutations at different time points across three dietary groups, we found that mutated SNVs in *P. distasonis* were particularly pronounced in the high-fiber diet group (HFID) (Fig. S5), further supporting the selective competition between BV9 and *P. distasonis*.

The results from pre- and post-coevolution *in vitro* coculture experiments complement the colonization patterns of BV9 under the three dietary interventions we designed: (1) Initially, there was no correlation between the original strains of BV9 and *P. distasonis* in either the normal or low-carb media (simulating a high-fat diet), which corresponds to BV9’s poor colonization in the ND and HFAD groups. (2) However, a competitive relationship emerged between BV9 and *P. distasonis* in the inulin-based medium. We hypothesize that BV9’s robust inulin metabolism capability is a key factor enabling its selective competition with *P. distasonis* and is a prerequisite for its successful colonization, as observed in the HFID group. (3) Finally, after coevolution within the host, a state of coexistence was achieved between the two bacteria. This was reflected in the absence of competitive interactions in the inulin medium during *in vitro* co-cultures of BV9 and *P. distasonis* strains isolated from the washout period.

The discovery of selective competition and the modifiability of bacterial competitive relationships offers new insights into probiotic colonization and even pathogen mitigation. For instance, enhancing probiotic colonization might be achieved by adding prebiotics and by targeting specific competitors, potentially using this phenomenon to deliberately alter the metabolism of pathogens if they are specific competitors of certain probiotics. Previous research has shown that selectively weakening particular bacteria is potentially feasible, e.g., bioengineered *Lactobacillus* probiotics (BLP) expressing *Listeria* adhesion protein (LAP) from a non-pathogenic *Listeria* competitively exclude pathogenic *Listeria* by occupying LAP receptors and heat shock protein 60, thereby reducing intestinal colonization and systemic spread^32^. Furthermore, given that competitive relationships can be modified and inherited, it may be necessary to consider domesticating a probiotic strain to enhance its colonization potential by adapting it through specific environmental or nutritional interventions. For example, previous studies have shown that the self-isolated strain is more effective in colonizing back to the mice, e.g., by exposing the candidate probiotic *E. coli* Nissle to the gastrointestinal tract of mice for several weeks under systematically altered dietary conditions, its adaptability was enhanced, leading to improved competitive ability in the gut and increased therapeutic efficacy in treating phenylketonuria^30^.

There are some limitations in the current study. For example, while the *in vitro* co-culture results support the variability and heritability of bacterial competitive relationships, our investigation was primarily at the genetic level. The potential mechanisms at other levels, such as epigenetics, transcriptomics, proteomics, and metabolomics, remain underexplored. In our future work, we plan to not only more systematically verify the enrichment differences at the genetic level in all genes related to inulin between the two bacteria but also incorporate methylomics and transcriptomics data to explore epigenetic differences in their methylation modifications. This approach could provide a more comprehensive explanation of their adaptive evolution. Additionally, we aim to further investigate the variability of the competitive relationships and delve deeper into their mechanisms, such as determining whether BV9 can colonize in the absence of *P. distasonis*, and what would happen to the gut microbiota if the inulin nutrient environment is altered after BV9 colonization—whether a new homeostatic state would form or if BV9 would be rapidly eliminated. Overall, this study, by revealing the specificity of niche competition during the colonization of exogenous bacteria, opens new avenues for modulating the gut microbiome.

## STAR METHODS

Detailed methods are provided in the online version of this paper and include the following:

### KEY RESOURCES TABLE

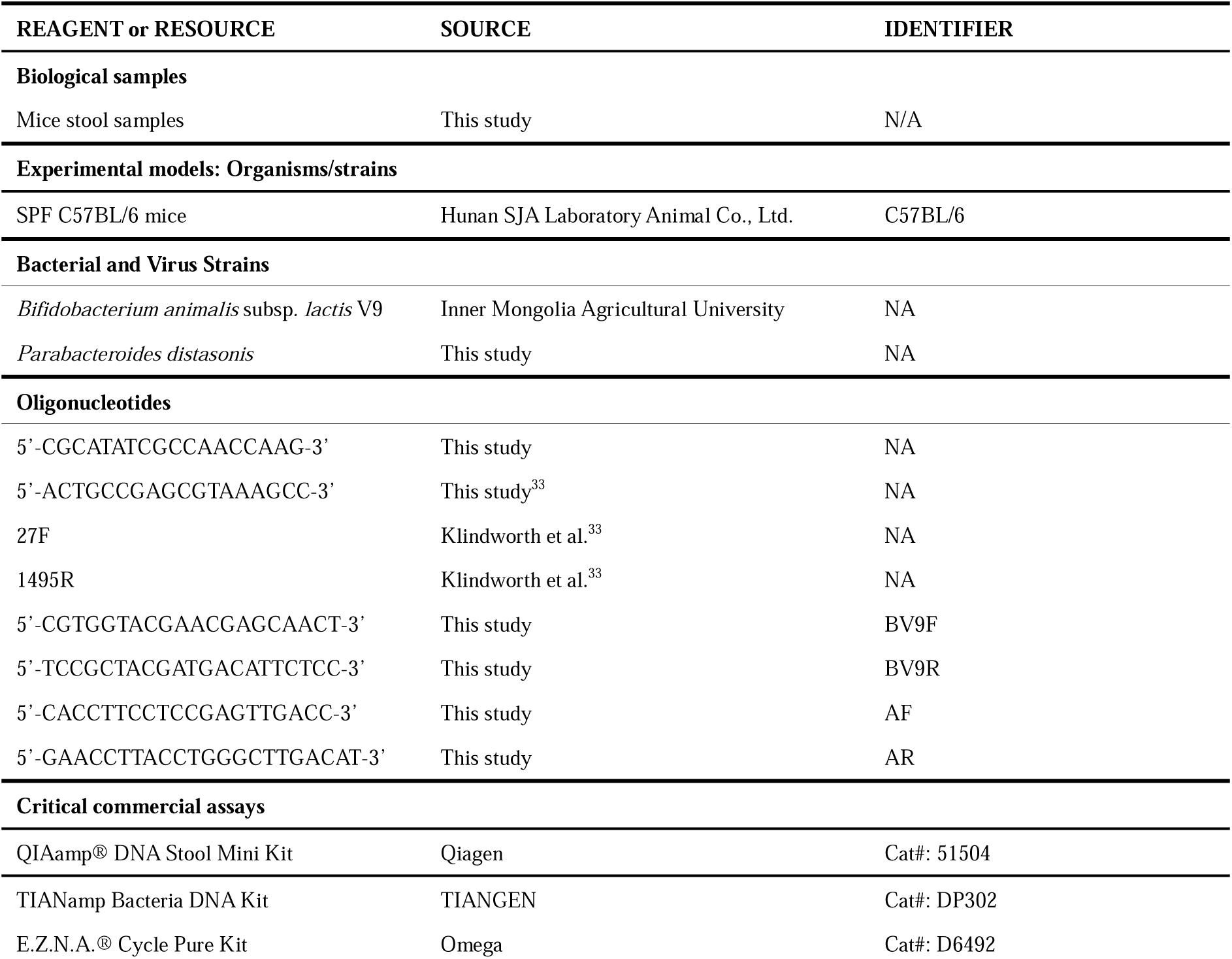
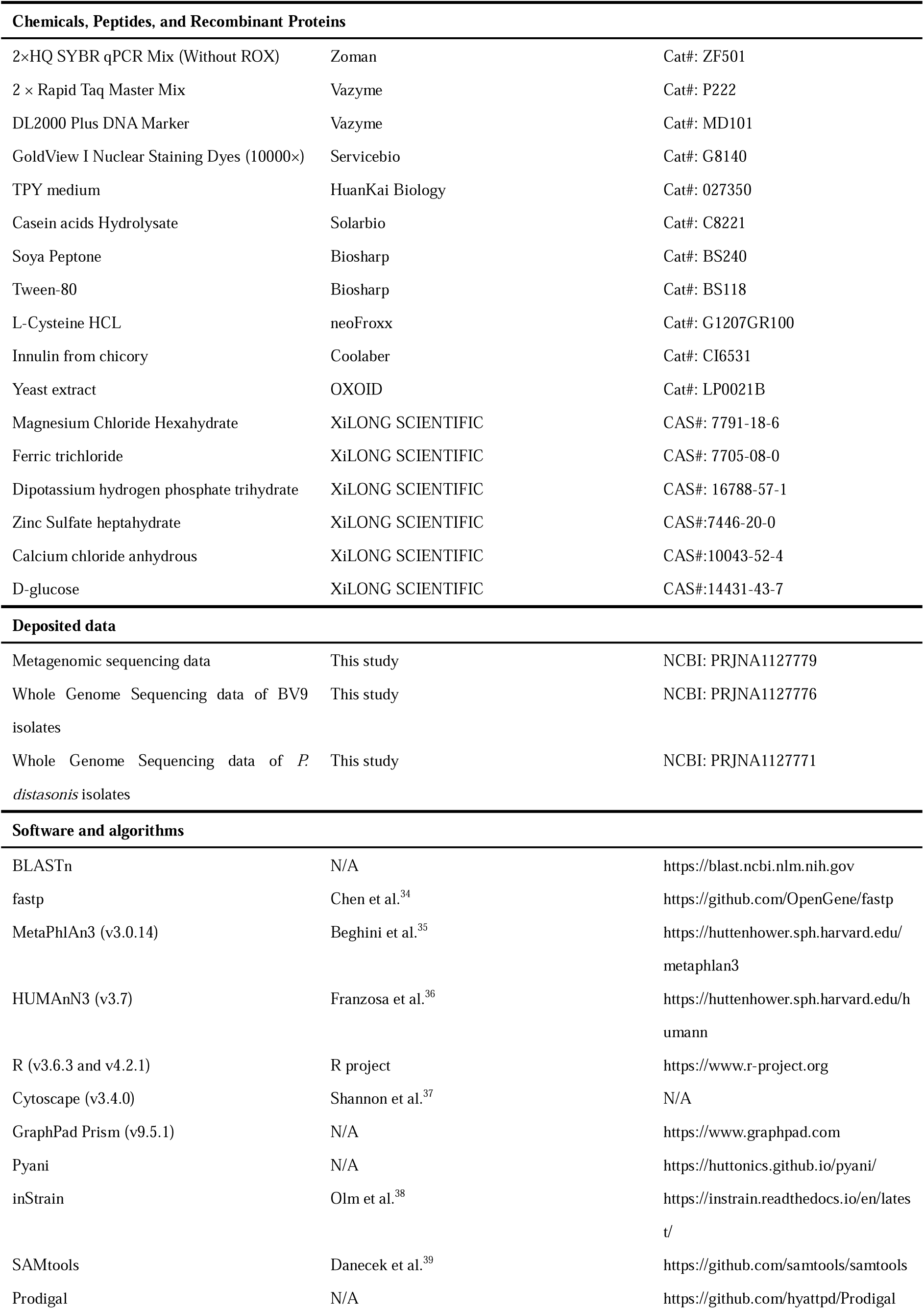
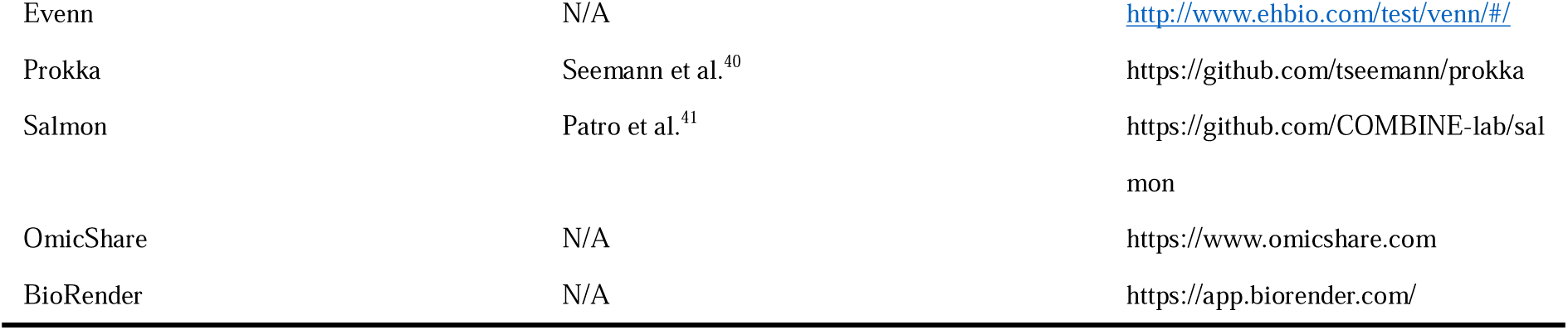

- RESOURCE AVAILABILITY
- Lead contact
- Further information and requests for resources and data should be directed to and will be fulfilled by the lead contact, Jiachao Zhang.
- Materials availability
- This study did not generate new unique reagents.
- Data and code availability
- The metagenomic sequence data and whole genome sequencing data have been deposited in the NCBI database (PRJNA1127779, PRJNA1127776 and PRJNA1127771).
- The scripts or codes used in this work have been deposited as in GitHub (https://github.com/hola1219/BV9).

## EXPERIMENTAL MODEL AND SUBJECTS RECRUITMENT

### Bacterial strain construction

#### Isolation of BV9 from Stool Samples

To evaluate the colonization efficiency of BV9, we isolated and cultured BV9 from the fecal samples collected daily. The isolation process is as follows: (1) Place the freshly collected fecal sample into a 10 mL centrifuge tube and add 1 mL of 0.85% sterile saline to homogenize; (2) To enrich BV9, transfer the entire homogenized mixture into 5 mL of fluid Trypticase Peptone Yeast Extract (TPY) medium, seal with sterile liquid paraffin, and incubate anaerobically for 12 hours; (3) After 12 hours, take 100 μL of the mixture and suspend it in 900 μL of sterile 0.85% saline for dilution to obtain dilutions of 10^4^, 10^5^, and 10^6^ for subsequent inoculation; (4) Take 100 μL of each dilution and spread it on solid TPY medium containing mupirocin at a final concentration of 75 μg/mL. Place the plates in an anaerobic jar with anaerobic gas-generating packs, label them with date and dilution, and incubate them anaerobically for 48 hours; (5) Finally, pick single colonies from the plates and amplify them in 5 mL of liquid TPY medium. After mixing, seal with 2 mL of sterile liquid paraffin and incubate anaerobically at 37°C for 48 hours for subsequent identification of BV9 mutants.

## METHOD DETAILS

### Dietary Patterns

We first sought to study the impact of various dietary patterns on the colonization of BV9. We focused on three representative dietary patterns: normal diet, high-fiber diet, and high-fat diet^42^. The normal diet, formulated in accordance with standard nutritional guidelines, was primarily composed of casein, corn starch, lard, minerals, and vitamins. This composition ensured a balanced intake of carbohydrates, proteins, and fats. Detailed information is provided in Table S1. The high-fiber diet was formulated by adding inulin to the normal diet to simulate high dietary fiber intake (37 g inulin/1000 kcal). The high-fat diet was formulated by increasing the lard content in the normal diet to simulate high-fat intake (60% kcal from fat). The detailed diet composition is listed in Table S1.

### Identification of BV9 in Isolations

BV9 is a *Bifidobacterium lactis* strain with excellent probiotic characteristics, isolated in 2005 from the intestinal tracts of a healthy Mongolian child in the grasslands of Inner Mongolia^43^. BV9 has been whole-genome sequenced^44^. To identify BV9 from all the isolates, DNA was first extracted and purified, then PCR with primers (forward: CGCATATCGCCAACCAAG; reverse: ACTGCCGAGCGTAAAGCC) was conducted for the first-round selection. Candidate isolates were then amplified using full-length bacterial identification primers (27F: GCAGAGTTCTCGGAGTCACGAAGAGTTTGATCCTGGCTCG; 1495R: AGCGGATCACTTCACACAGGACTACGGCTACCTTGTTACGA)^33^. 16S rRNA sequencing was performed by Qingdao Pengxiang Biotechnology Co., Ltd. Finally, the identification and confirmation of BV9 in candidate isolates were performed using NCBI’s BLASTn (https://blast.ncbi.nlm.nih.gov/blast.cgi) with identify higher than 99%.

### Whole Metagenome Sequencing and Analysis

Fecal samples collected on Days 1, 7, 14, 21, and 28 were first labeled and stored at −80°C and then sent for the whole metagenome sequencing (WMS). The QIAamp DNA Stool Mini Kit was used to isolate and purify DNA from mouse fecal samples. We used 0.8% agarose gel electrophoresis to perform preliminary quality analysis on the purified DNA, then used Nanodrop to detect the purity of the sample DNA by OD260/280 value, and finally quantified the DNA concentration using qubit2.0. WMS was performed using Illumina HiSeq 2500 sequencer at Beijing Novogene Technology Co., Ltd., and a DNA library of paired-end reads (2×150 bp) was generated before sequencing. The raw data were preprocessed using fastp^34^ (https://github.com/OpenGene/fastp) to remove adapter sequences and filter out the following paired reads: (1) reads with low-quality bases (Q ≤ 5) exceeding 50% of the total read length and (2) reads with N content exceeding 10% of the total read length. After quality control, the cleaned raw data were aligned to the host genome to filter out reads potentially originating from the host. The average sequencing depth for each sample was 6X.

To generate the taxonomic and functional profiling results for the WMS data, we first applied MetaPhlAn3 to identify the microbial species and estimate their relative abundances^35^. Then, based on the UniRef90 database, we employed HUMAnN3 to decode the functional profiles of the gut microbiome^45^.

### Whole Genome Sequencing of BV9 and P. distasonis

Whole genome sequencing (WGS) for all BV9 and *P. distasonis* isolates (confirmed by 16S rRNA sequencing) was performed on the Illumina Hiseq 2500 platform at Beijing Novogene Technology Co., Ltd., using paired-end (2×150 bp) sequencing. To ensure the accuracy and reliability of subsequent analyses, quality control was performed on the raw sequencing data.

The raw data were filtered using fastp^34^ to obtain clean data. The specific parameters used were: --in1 input_1.fq.gz, --out1 out_1.fq.gz, --in2 input_2.fq.gz, --out2 out_2.fq.gz, -g, -q 5, -u 50, -n 15, -l 150, --min_trim_length 10, --overlap_diff_limit 1, --overlap_diff_percent_limit 10, -j output.json, -h output.html. The average sequencing depth for each sample was 350X.

### Construction of Co-occurrence Networks

To understand the potential interactions between BV9 and native gut microbiota under different dietary patterns, we aggregated species-level profiles generated by the WMS data on days 1, 7, 14, 21, and 28 from each group (i.e., ND, HFID, and HFAD) to construct the group-specific co-occurrence network (Fig. 3a). Co-occurrence networks were constructed using R software (with WGCNA packages; Spearman test) and visualized with Cytoscape 3.4.0. We also visualized the associations between BV9 and other bacteria in different diet groups using the “pheatmap” package (Fig. 3b). Moreover, line plots created with GraphPad Prism (Version 9.5.1) illustrated the relative abundance changes of BV9 and *P. distasonis* in three groups, and the T-test was performed, with “*” indicating p < 0.05 (Fig. 3c). Additionally, the “ggplot2” and “ggpubr” packages were used to calculate and visualize the correlation between the two species (Spearman test) (Fig. 3d).

### Comparative Genomic and Functional Analysis

To explore the selectivity of BV9 against its competitors, we conducted a comparative analysis of the genomic structure and functional capabilities of BV9, its related species, and the top 20 species by relative abundance. In terms of genomic structure, we first downloaded the reference sequences for these species from the NCBI database. Subsequently, we used Pyani to calculate the Average Nucleotide Identity (ANI) values between BV9 and these species. For functional capabilities, we performed functional annotation of these genomes using the KEGG Automatic Annotation Server^46^ (www.genome.jp/tools/kaas/), followed by further analysis of the annotated results with the Mapper tool in the Kyoto Encyclopedia of Genes and Genomes (www.genome.jp/kegg/mapper/). The results were plotted using GraphPad Prism (Version 9.5.1, Fig. 3e).

### SNVs Calling

To investigate the genetic variations in BV9 and *P.distasonis* isolates, we used inStrain software (v1.0.0) to perform single nucleotide variants (SNVs) calling by using the WGS data of the isolates^38^. First, based on the sequence information of the reference strains, we used Bowtie2 to create the library and SAMtools to convert the library to BAM format^47,48^. Next, we used Prodigal to annotate the reference strains, obtaining their functional genomes. Finally, we used inStrain to analyze SNVs information in each isolate (inStrain profile sample.bam ref.fna -c 100 -f 0.49 -o sample.profile -p 24 -g ref_genes.fna). On average, each sample yielded 35,026 SNVs.

### Candidate SNVs Selection and Pathway Enrichment Analysis

To elucidate the mechanisms of competition and co-evolution between BV9 and *P. distasonis*, we used Evenn to analyze SNVs at different time points and obtained candidate SNVs (http://www.ehbio.com/test/venn/<u>#/</u>). Venn diagrams were generated using Evenn (Fig. 4d, e, f). We then classified these candidate SNVs into two categories: coding and non-coding regions. For SNVs located in coding regions, we evaluated whether they were non-synonymous mutations and if the genes containing them were under positive selection. For SNVs in non-coding regions, we performed differential abundance analysis and differential enrichment analysis on the downstream genes and pathways they influence. These analyses were conducted for both BV9 and *P. distasonis* as follows: To harmonize the gene annotations, we first re-annotated the genomes of BV9 and *P. distasonis* reference strains using Prokka^40^. Next, Salmon (1.10.2) was employed to quantify the sequencing depth of genes using the WGS data of all isolates^41^ (salmon quant -i sample -l A −1 sample_1.fastq.gz −2 sample_2.fastq.gz -o sample.quant). Differential abundance analysis was then conducted using the “edgeR” package in R to identify differentially abundant genes among the HFID, HFAD, and ND groups. Finally, differential enrichment analysis for pathways, including the creation of bubble diagrams for enriched KEGG pathways, was performed using OmicShare (https://www.omicshare.com) with default parameters.

To explore the co-evolutionary mechanism between BV9 and *P. distasonis* at the genetic level, we applied a log2 fold change threshold of 0.4 to further refine the differential genes and visualized the results using the “ggplot2” package (Fig. 4g, 4h).

### Co-culture Experiments and Quantitative Real-time PCR

To explore the relationship between BV9 and *P. distasonis*, we co-cultured the original BV9 and *P. distasonis* as well as the isolated BV9 and *P. distasonis* at Day 3 during the washout period. To simulate the different nutritional environments, we modified the medium composition accordingly. We used: (1) standard fluid TPY medium (without additional nutrients) to simulate normal dietary conditions (ND); (2) we added inulin to the TPY medium to simulate a high-fiber diet (HFID); and (3) we reduced the carbon content in TPY medium to simulate a high-fat diet (HFAD). BV9 and *P. distasonis* were inoculated in equal amounts into the three media types and co-cultured for 48 hours. Samples were collected every four hours, with 1 mL (stored at −80°C) collected for each time point. After collecting samples from all time points, DNA was extracted and purified as described earlier. Subsequently, quantitative real-time PCR (qPCR) was performed with primer (BV9F: CGTGGTACGAACGAGCAACT, BV9R: TCCGCTACGATGACATTCTCC; AF: CACCTTCCTCCGAGTTGACC, AR: GAACCTTACCTGGGCTTGACAT). Specifically, the reaction parameters were as follows: 94°C pre-denaturation for five minutes, 35 cycles of 94°C denaturation for 30 seconds, 55°C annealing for 45 seconds, 72°C extension for 40 seconds, and a final extension at 72°C for ten minutes. qPCR data were statistically analyzed using the “ggpubr” package in R and visualized by using “ggplot2” package (Fig.6).

## QUANTIFICATION AND STATISTICAL ANALYSIS

### Statistical Analysis

All schematic figures and flow charts were created by using BioRender (https://app.biorender.com/) (Fig. 1, Fig. 4a, and Fig. 5). To accurately describe the colonization of probiotics in different groups, we used Prism 9 to statistically analyze and visualize the number of isolated strains in the ND, HFID, and HFAD groups (Fig. 2a). Additionally, for the WMS data, we utilized the “ggplot2” and “ggbeeswarm” packages in R to create box plots showing the relative abundance of BV9 (Fig. 2b), to double-check the colonization status of the microbial community *in vivo*.

To comprehensively describe the dynamic changes in gut microbiota following the probiotic intervention, we used the R package “vegan” to perform the alpha diversity analysis (Shannon index) and Principal Coordinate Analysis (PCoA) for taxonomic profiles generated from WMS data. Loess-fitting curve and scatter plots were created using the “ggplot2” package (Fig. 2c, d). Ridge plots based on the Bray-Curtis dissimilarity were generated using the “ggridges” package (Fig. 2e). The Kruskal-Wallis test and ALDEx2 were used to identify differential species between groups, and the results were visualized using the “pheatmap” package (Fig. S2a). Differential pathways were calculated and screened using the “DESeq2” package in R, with visualization also performed using the “pheatmap” package (Fig.S2b).

## Acknowledgments

This research was supported by the National Natural Science Foundation of China (32222066).

## Author contributions

JZ, YL, ZS and ZH conceived the concept and designed the study. ZH, QZ, LD, DZ XL and SJ performed all experiments and prepared the samples. ZH and ZS carried out the metagenome analyses with significant contributions from YL. ZS and ZH wrote the paper. YL, ZS and JZ provided advice and revised the manuscript extensively.

