## Supplementary figures and images for "Competition and Compromise between Exogenous Probiotics and Native Microbiota"

### Supplemental Figure1

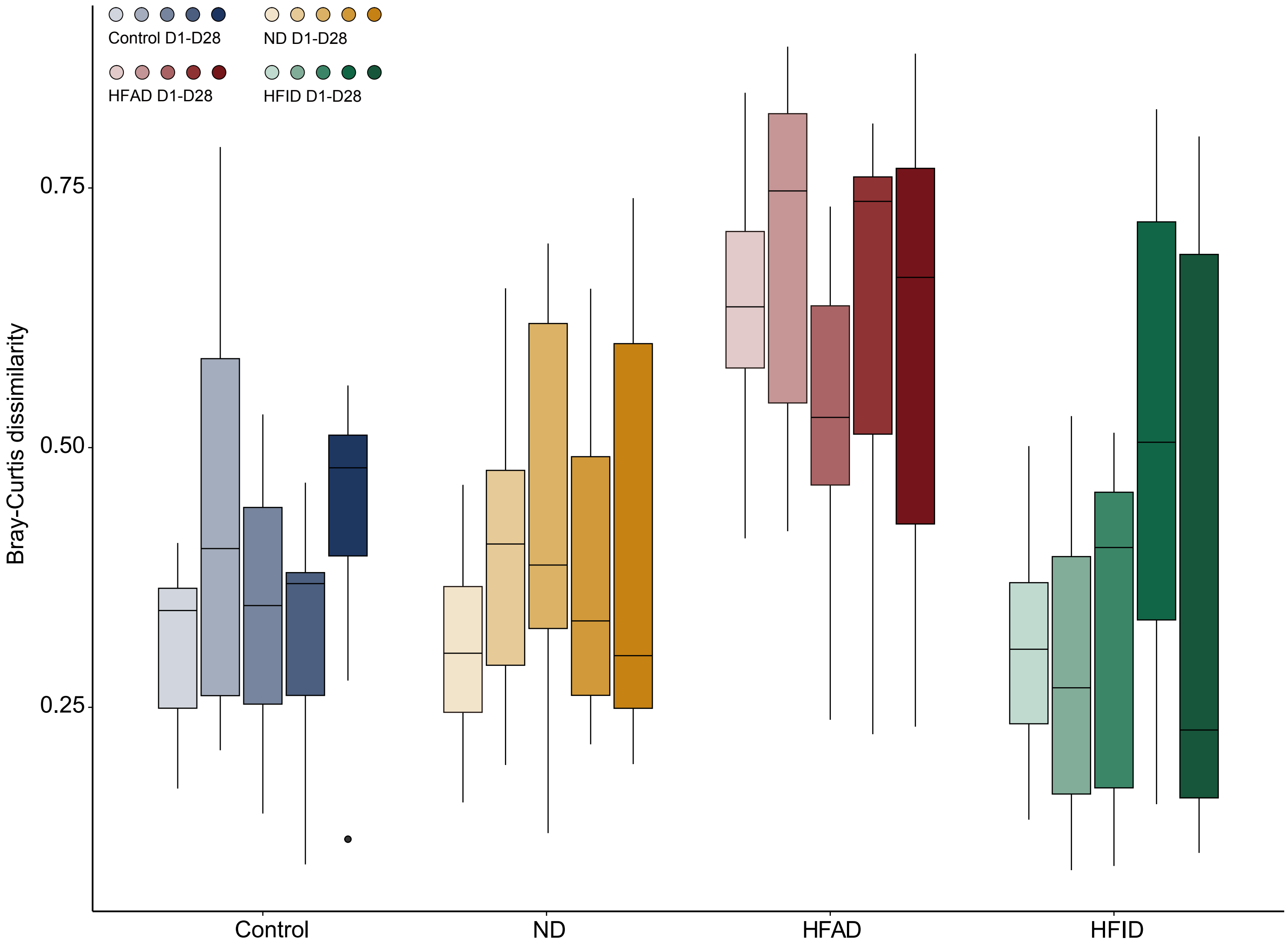

### Supplemental Figure2

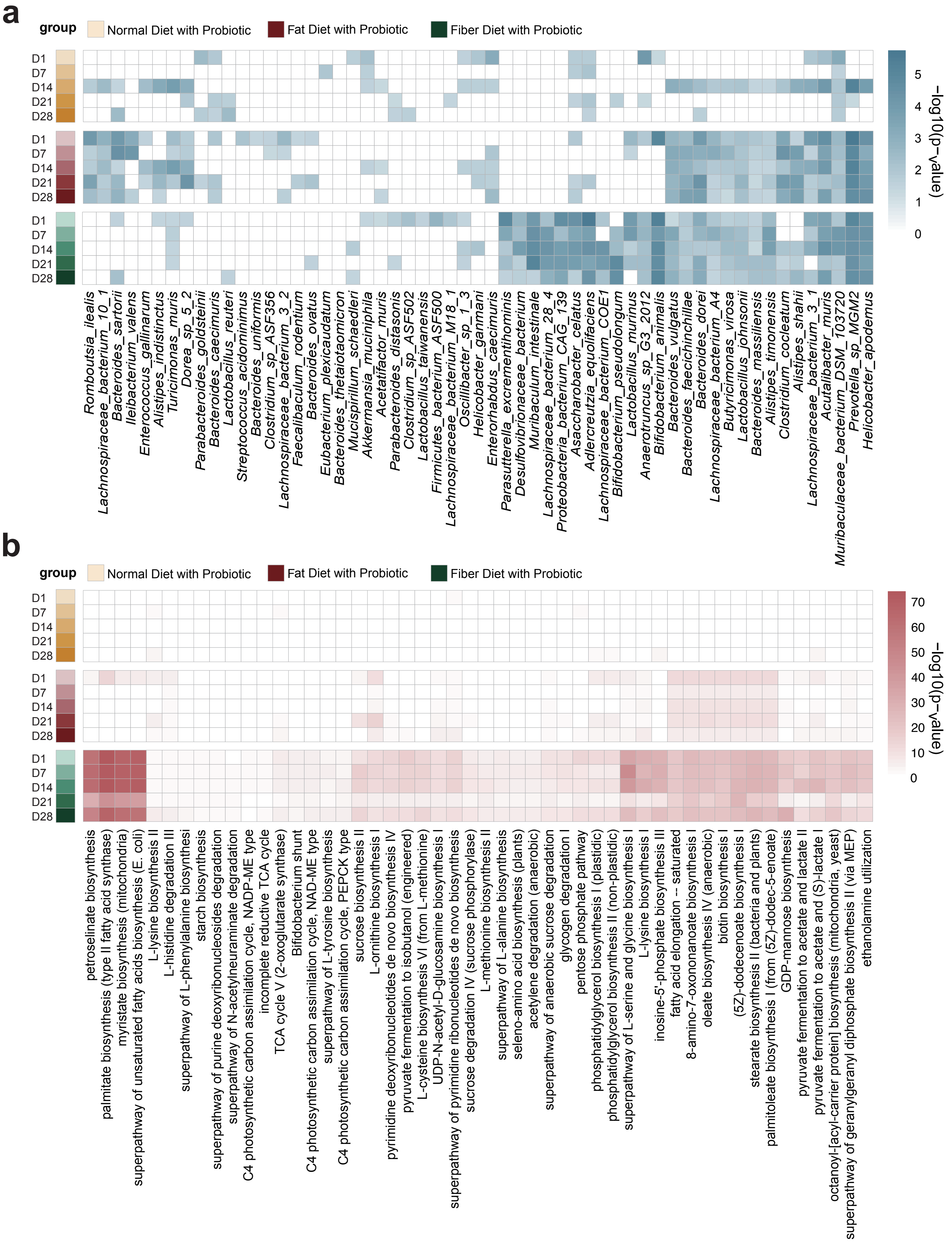

### Supplemental Figure3

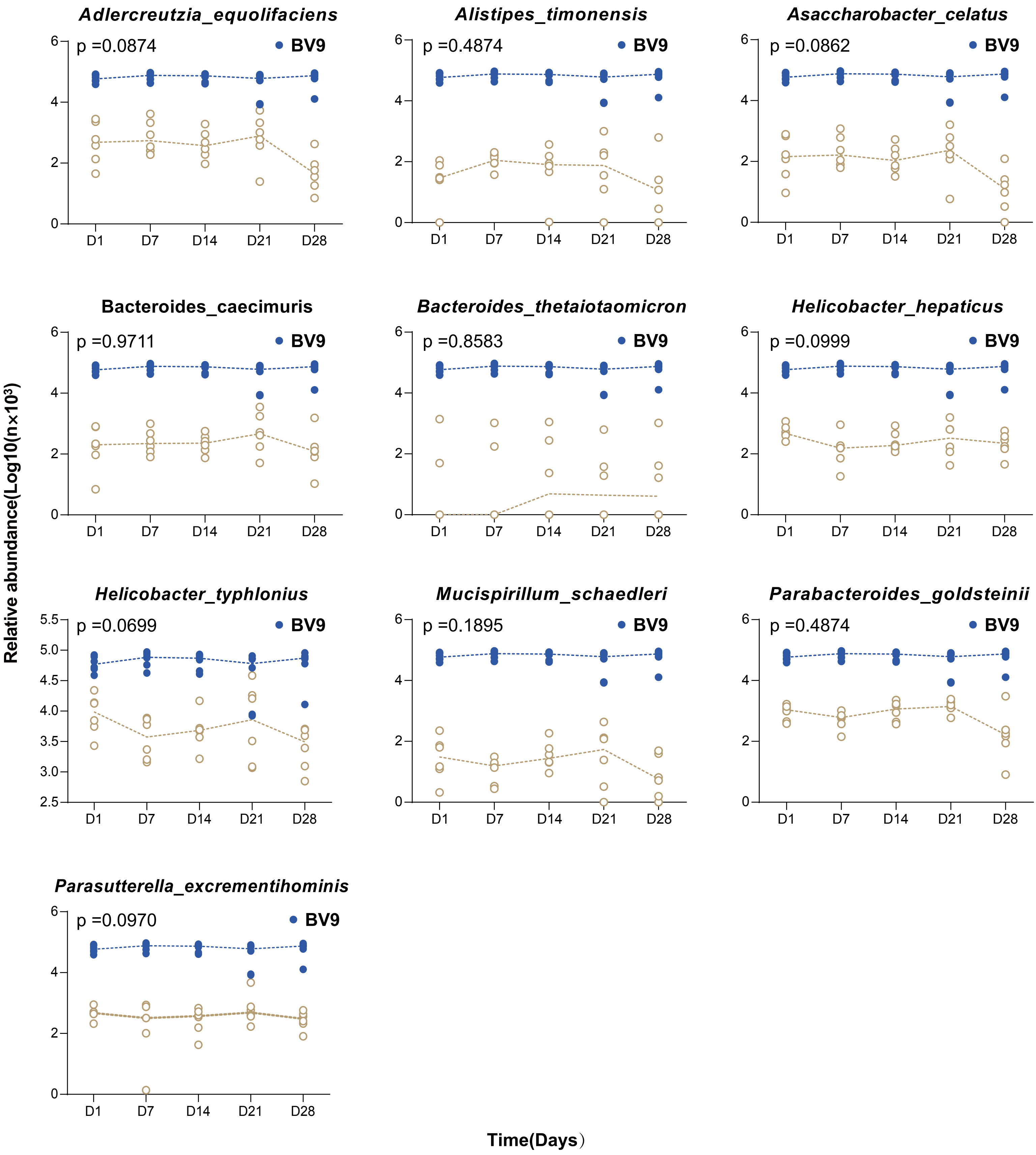

### Supplemental Figure4

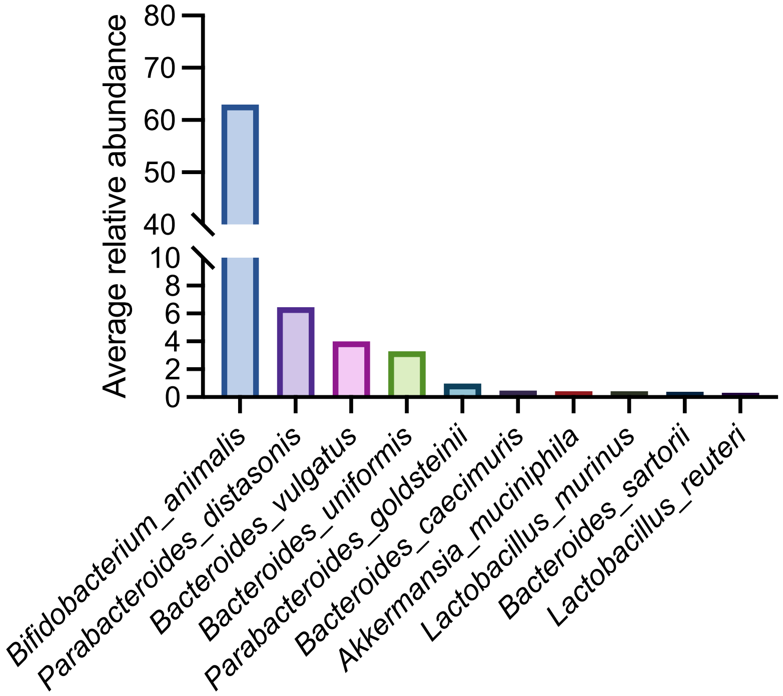
