## Supplemental Figure5 for "Competition and Compromise between Exogenous Probiotics and Native Microbiota"

| | ND | | | | | HFAD | | | | | HFID | | | | |
|---|---|---|---|---|---|---|---|---|---|---|---|---|---|---|---|
| | D1 | D7 | D14 | D21 | D28 | D1 | D7 | D14 | D21 | D28 | D1 | D7 | D14 | D21 | D28 |
| *Bifidobacterium animals* | | | 518 | 21 | 364 | 7 | 28 | 14 | 7 | 18 | 34136 | 37386 | 36582 | 26991 | 42573 |
| *Acutalibacter muris* | | 1 | 1 | | | 17 | | | 19 | 10 | | | | | |
| *Alistipes shahii* | | 4 | | | | | | | | | | | 2 | | |
| *Bacteroides caecimuris* | 5 | 86 | | | | 11 | | 1 | 1 | 2 | 4 | 2 | 1 | 1 | 1 |
| *Bacteroides thetaiotaomicron* | | | | | | | | | | | | | | | |
| *Parabacteroides distasonis* | | 1 | | | | 8 | 1 | 1 | | 1 | 10 | 36 | 44 | 11 | 8 |
| *Parabacteroides goldsteinii* | | | | | | | | | | | | 6 | 5 | | |
| *Parasutterella excrementihominis* | | | | 1 | | 9 | 8 | 12 | 10 | 8 | 6 | 6 | 7 | | |

Median # of SNPs: 0, 10000, 20000, 30000, 40000
